# Batch Effect Removal via Batch-Free Encoding

**DOI:** 10.1101/380816

**Authors:** Uri Shaham

## Abstract

Biological measurements often contain systematic errors, also known as “batch effects”, which may invalidate downstream analysis when not handled correctly. The problem of removing batch effects is of major importance in the biological community. Despite recent advances in this direction via deep learning techniques, most current methods may not fully preserve the true biological patterns the data contains. In this work we propose a deep learning approach for batch effect removal. The crux of our approach is learning a batch-free encoding of the data, representing its intrinsic biological properties, but not batch effects. In addition, we also encode the systematic factors through a decoding mechanism and require accurate reconstruction of the data. Altogether, this allows us to fully preserve the true biological patterns represented in the data. Experimental results are reported on data obtained from two high throughput technologies, mass cytometry and single-cell RNA-seq. Beyond good performance on training data, we also observe that our system performs well on test data obtained from new patients, which was not available at training time. Our method is easy to handle, a publicly available code can be found at https://github.com/ushaham/BatchEffectRemoval2018.

## 1 Introduction

Biological measurements are typically affected by systematic errors, i.e., factors which depend on experimental conditions but not on the phenomena being measured. Such factors are known in the biological community as *batch effects*. As the magnitude of batch effects can potentially be significant comparing to the magnitude of the biological signal of interest, ignoring the existence of batch effects may lead to false results invalidating a downstream analysis. For this reason, the problem of batch effect removal has drawn major interest in the biological community [11].

Obviously, the specific effect of the experimental conditions on the measurements is never known. However, it is usually easy to obtain two or more control samples, where each sample is being collected at one set of experimental conditions (i.e., a batch), and both samples correspond to the same underlying biological state. This setting naturally promotes machine learning techniques, that learn patterns from examples, to use for batch effect removal. Two important challenges for any such machine learning approach for batch effect removal are (i) its ability to generalize, i.e., to remove batch effects from samples which differ in distribution than the ones used for training and (ii) not distorting true biological signal represented in the data. Obtaining good performance on arbitrary test data that differ in distribution from the training data is a fundamental difficulty for any machine learning algorithm, and good generalization ability can thus never be guaranteed in principle, for any batch effect removal procedure. Yet, experimental results can be used to evaluate the generalization ability of proposed procedures. Preservation of biological signal while removing batch effects is crucial, as otherwise, calibrated data does not fully reflect the true biological state of a subject, which might invalidate any subsequent downstream analysis as well.

In last decade deep learning have achieved tremendous performance on various machine learning tasks across numerous domains, and several recent works have utilized it for batch effect removal. Despite making important progress in this direction, most recently proposed deep learning methods for batch effect removal have fundamental weaknesses preventing them from successfully fulfilling the second challenge above.

In this work, given biological measurement data, we seek to obtain a representation (or a code) of the data which encodes solely biological signal but not batch effects. In addition, we also require that our code, along with a decoding mechanism will allow reconstruction of the data, so that the entire true biological signal will not be lost or distorted. The approach presented in this manuscript contains an encoder-decoder deep learning system in which (i) batch effects are stripped off raw measurement data via an encoding mechanism, (ii) the data is represented in a code space, in which the code corresponds to its underlying biological state and (iii) the measurement data is reconstructed by a decoding mechanism in which the batch effects are hard-coded. In order to obtain such desirable code we utilize adversarial loss, a powerful deep learning technique. Requiring invariance of the encoding to batch effects, along with good reconstruction performance encourages our system to fully preserve biological properties represented in the data. We demonstrate the performance of our system on two novel biological techniques, mass cytometry and single-cell RNA sequencing (scRNA-seq). Along with obtaining good performance in removal of batch effect from the training samples, we observe that the proposed system performs well also at test time, on samples arising from new distributions, which were unknown during training. This manuscript is accompanied with an easy to use and publicly available Python code.

The remainder of this manuscript is organized as follows. In Section 2 we review several recent deep learning-based approaches for batch effect removal, as well as related works from other domains. In Section 3 we describe our proposed approach. Experimental results are reported in Section 4. Section 5 briefly concludes the manuscript.

## 2 Related work

Several research groups recently proposed deep learning-based approaches for batch effect removal. The Kluger research group proposed an approach based on networks trained to minimize maximum mean discrepancy (MMD, [6]) distance between a source and a target distribution [17, 12]. As calibration by minimizing the distribution distance alone can distort important biological properties of the data, they used residual nets [8], where the learned calibration map is encouraged to be of small magnitude, implicitly preserving biological characteristics of the data. In this manuscript we take a different path to ensure that biological properties are preserved, by explicitly requiring the encoding to be batch-free. In addition, while they demonstrated impressive experimental results on training data, the ability of such approach to perform well on test data from arbitrary distributions is questionable.

Using MMD for batch correction was later used also by the Krishnaswamy research group in [3]. A second work by this group [1] used generative adversarial nets [5] rather than MMD loss to match the distributions. In addition, this approach requires labeled subsets of the data and applies a mechanism that encourages labeled subpopulations of the data to maintain pairwise correspondence, in order not to distort biological structures. Beyond the fact that this approach is no longer unsupervised and requires domain knowledge, the amount of labeled data that might be needed to achieve reasonable performance can unfortunately be large, for example in case of scRNA-seq data, which typically consists of many clusters. In contrast, the approach presented here is purely unsupervised and does not rely on any domain knowledge or label information.

A recent work by the Krishnaswamy group [2] proposed to address the generalization issue by learning new representation for the data, identifying dimensions which correspond to batch differences (while those which do not may correspond to biological properties), and aligning the distributions of the batches for each such “batch dimension”. This approach is based on two powerful assumptions. First, they assume that there exist dimensions in this representation that encode only batch differences but not biological properties, so that the batch differences can be corrected by aligning the two batches in these dimensions. Second, the alignment is being done for each dimension separately, which implicitly assumes that these dimensions are uncorrelated. Both assumptions are unrealistic in the general case, as no elements of their algorithm encourage the learned representation to satisfy them. Moreover, even if such “batch dimensions” and “biology dimensions” do exist in the encoding of the control samples, there is no particular reason to assume that they will be consistent with the representations of arbitrary samples arriving at test time.

Our approach is related to that of [13] and the more recent work of [16], who propose domain adaptation techniques based on learning domain-invariant features, where the first one utilizes MMD while the second one utilizes adversarial training. It is also closely related to [15], where domain-invariant representation is learned for music style transfer. Finally, domain invariant representation is learned also in [19], utilizing Wasserstein distance, which we use as well. They train a model to “forget” specifically chosen domain information. However, for batch effect removal, their method may “forget” also true biological signal. In contrast, we do not forget any information in the data, as we require reconstruction of the data. Rather, we split the information to biological information, which is present in the code, and batch information, which is hard coded into the decoding mechanism.

## 3 Methods

### 3.1 Problem setup

Let 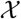 be an arbitrary space, corresponding to the collection of possible intrinsic biological states of a subject. Let 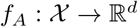 and 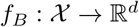 be two measuring environments (i.e., two instruments, or two sets of laboratory conditions), creating two batches of measurements. Let *p* be a distribution on 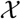, which corresponds to a specific biological state of a subject and let *X*_1_, *X*_2_ be two unobserved sets of iid samples from *p*, of arbitrary sizes *n*_1_ and *n*_2_, respectively.

We observe two batches of measurement data *X_A_* = *f_A_*(*X*_1_) and *X_B_* = *f_B_* (*X*_2_). Our goal is to use *X_A_* and *X_B_* to learn a system that “calibrates” batch differences, i.e., we would like a calibration system that when trained on *X_A_* and *X_B_*, will output two samples *X′_A_*, *X′_B_* having the same distribution and same underlying biological properties.

In addition, a desirable property of such system is its ability to perform the calibration well also on new test samples. More formally, consider two future observations 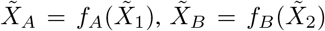, where 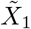 and 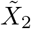 are sampled from unknown distributions *q*_1_ and *q*_2_, respectively (thus corresponding to samples with possibly different biological conditions, which may also differ from the training distribution *p*). We would like our system to enable us to compare the underlying intrinsic biological states represented in 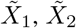 in a way that would not be affected by the differences between *f_A_* and *f_B_*. As explained above, such property, while desired, can not be guaranteed in principle for any batch effect removal technique.

### 3.2 System design

#### 3.2.1 Rational

Our system consists of a shared encoder *E* and two decoders *D_A_*, *D_B_* which reconstruct the data for each batch. Reconstruction of the data requires that no information, and specifically information on the internal biological state of the subject, is lost. In addition, we train a discriminator network Disc and use adversarial loss to obtain an encoding in which the distributions of two control samples, corresponding to the same intrinsic biological state, cannot be distinguished, despite the fact that the two samples were collected in different batches. Together, this implies that while the code is batch-free, all batch information is hard coded into the decoders. Hence, intuitively, one may think of our proposed system as an encoder which removes the batch information from the measurements and encodes the data in a batch-free space, and decoders that given such code, put the batch information back on to reconstruct the data. The two conditions we require, i.e., (i) batch-free encoding and (ii) reconstruction, imply together that the code has to contain complete information on the underlying biological state represented in the data, as otherwise, a good reconstruction will not be possible. Below we give a short technical background of each component and describe the specific setup of our proposed system.

#### 3.2.2 Variational autoencoder

We implement each of (*E*, *D_A_*) and (*E*, *D_B_*) as a variational autoencoder (VAE, [10]). A variational autoencoder is a pair (*E*, *D*) of networks, where the encoder E maps each data point 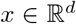 to a distribution *E*(*x*) over the code space. We follow a typical setting where *E*(*x*) is a *r*-dimensional Gaussian with a diagonal covariance matrix, parametrized by its mean vector and the covariance diagonal. The decoder *D* maps a code *z* back to a distribution *D*(·|*z*) over the input space. A VAE is trained to maximize a lower bound on the likelihood of its training data. Specifically, for data coming from a training distribution 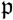, a VAE is trained to minimize the loss

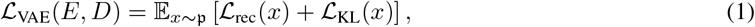

where

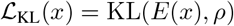

is a reconstruction term and

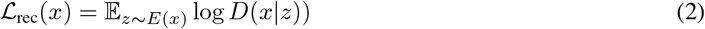

is a term penalizing the distance of the code distribution from a generic standard distribution *ρ* (usually a standard *r*-dimensional Gaussian). Typically, we consider the decoder output *D*(·|*z*) to be the mean of a d-dimensional Gaussian with identity covariance, in which case the reconstruction loss (2) is proportional to the squared error

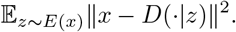

In our experiments, we used a more flexible variant of VAE [9], in which the KL term in (1) is multiplied by a tunable parameter *β*, so that the VAE loss becomes

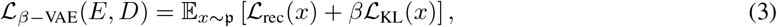

#### 3.2.3 Adversarial training

We would like the encoding of the data to be batch-free, i.e., to correspond solely to the intrinsic biological state the data represents but not to the measuring environment. Observe that the training data *X_A_* and *X_B_* correspond to the same intrinsic biological state represented by *p*. Denote by *p_A_* the distribution of *E*(*f_A_*(*X*)), where *X* ~ *p*, and similarly *p*_B_. In a code space which encodes solely the underlying biological state, the distributions *p_A_* and *p_B_* should be similar. To achieve this, we use adversarial framework, a powerful technique in deep learning [5], with which we minimize the Wasserstein distance between *p_A_* and *p_B_*, defined (via the Kantorovich-Rubinstein duality) as

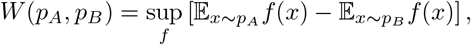

where the supremum is taken over all 1-Lipschitz functions. Specifically, we train a discriminator network Disc: 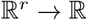, implemented as a Wasserstein GAN [4] discriminator. The discriminator approximates the Wasserstein distance between *p_A_* and *p_B_*. We then require the encoder *E* to maximize the discriminator loss, by producing code distributions with small Wasserstein distance. We train the discriminator to minimize the loss

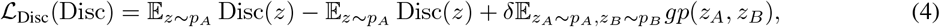

where *δ* is a user-specified small positive constant and *gp* is the gradient penalty loss introduced in [7], which controls the Lipschitz constant of the discriminator via penalizing its gradient wrt its input, i.e.,

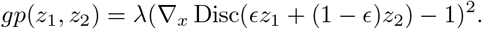

where *ϵ* is a random *U*(0, 1) sample.

The Encoder *E* is trained to minimize

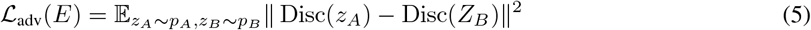

#### 3.2.4 Putting it together

Altogether, we train (*E*, *D_A_*, *D_B_*) and Disc in parallel, where the Disc is trained to minimize (4), while (*E*, *D_A_*, *D_B_*) are trained to minimize

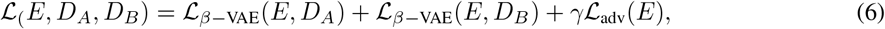

where γ is another user-specified small positive constant. The batch-free encoding allows for calibration of the data and representing it in the input space: once the system is trained, one encodes data from two batches and reconstruct the data from both batches using the same decoder.

## 4 Experiments

### 4.1 CyTOF data

CyTOF [18] is a mass cytometry technology that allows simultaneous measurements of multiple protein markers in each cell of a specimen (e.g. a blood sample), typically consisting of 10^3^-10^6^ cells.

We perform our experiments on a subset of the publicly available data used in [17]. The data contains four samples, belonging to two patients, where each patient has two samples collected on different days on the same CyTOF machine. We arbitrarily choose patient 2 for training and patient 1 for testing, where for each patient, we consider samples from different days as different batches. All samples had dimension d = 25 and contained 1800-5000 cells each. For a full description of the data and a specification of the markers we refer the reader to Section 4.2.1 and the Supplementary Section S1 in [17]. The data was pre-processed using a log transform *x* ← log(1 + *x*). Unlike [17] we did not use any denoising autoencoders to remove zero entries from the data.

Figure 1 shows the reconstructions of the train and test data, projected onto the subspace of first two principal components of the train data. For each batch, the reconstruction was obtained by encoding its data and decoding it via the corresponding decoder. Figure 2 shows the calibration of train and test data. The calibration was done by encoding the data and using the target batch decoder to decode both batches. As can be seen in both figures, the quality of the reconstruction and calibration of the test data (i.e., data from patient 1) is similar to that of the train data (i.e., data from patient 2).

**Figure 1:**
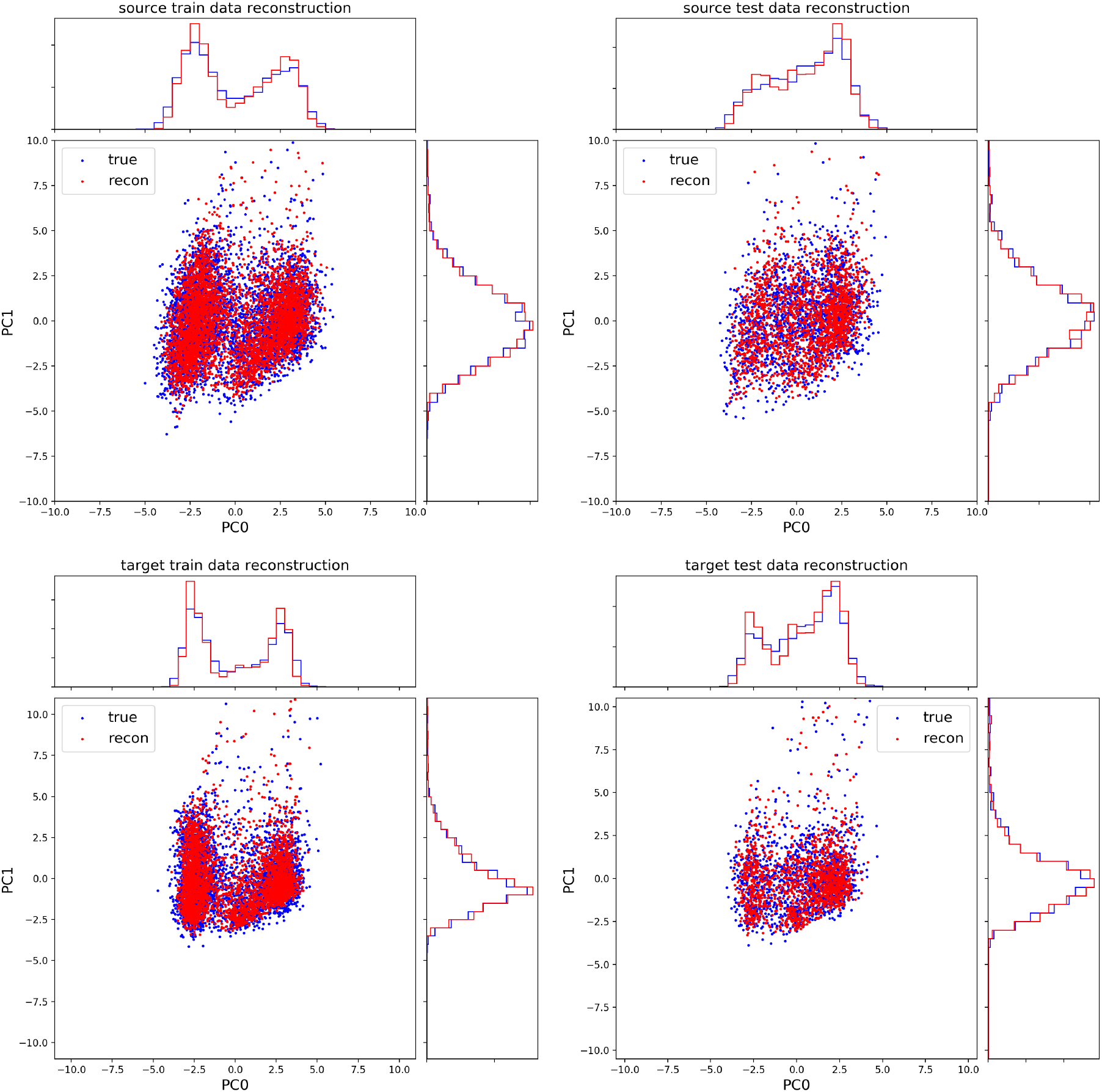
Reconstructions of cytof data. Data from patient 2 was used for training (left) and from patient 1 for testing (right). The terms “source” and “target” refer to the two batches (top and bottom, respectively). In all plots the inputs to the encoder are marked in blue, and the outputs from the corresponding decoder are marked in red.

**Figure 2:**
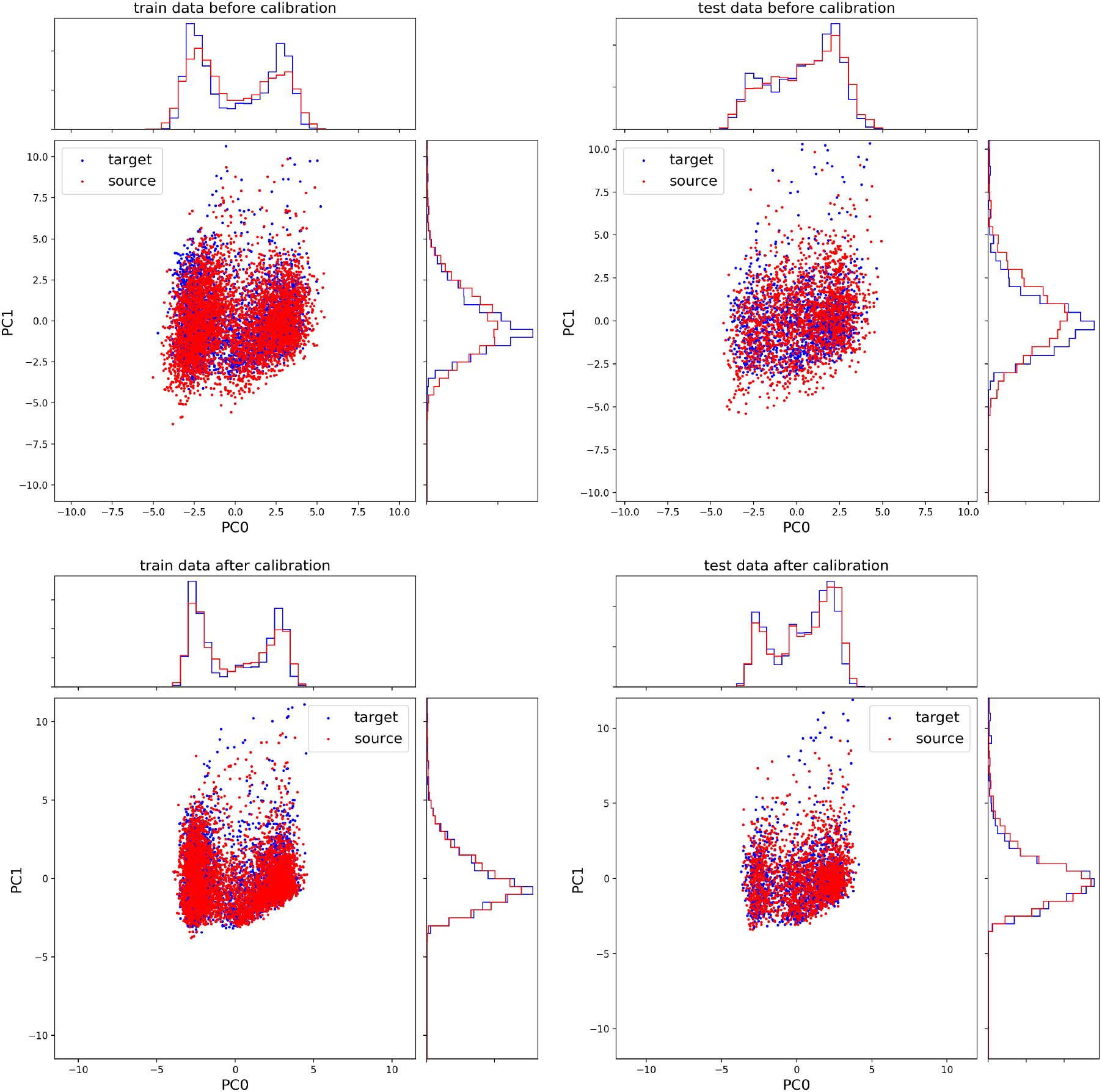
Calibration of cytof data. Data from patient 2 was used for training (left) and from patient 1 for testing (right). Top: data before calibration. Bottom: Data after calibration. In all plots “source” and “target” refer to the two batches.

Figure 3 shows the calibration of test data for each of three individual markers. The results are similar to those on all other 22 markers as well. As can be seen, the cumulative distributions of each marker are significantly closer after calibration than before calibration.

**Figure 3:**
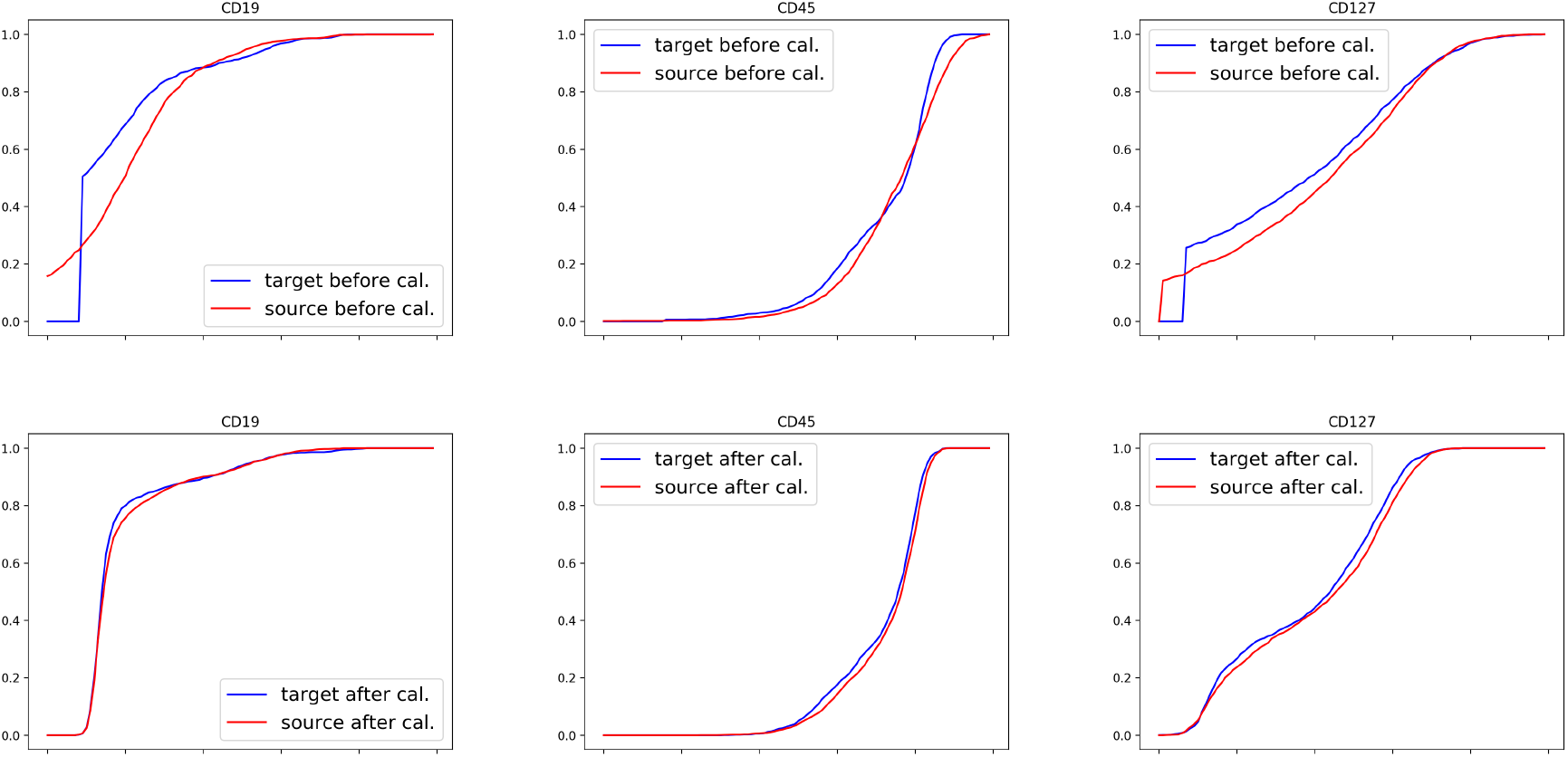
Calibration cytof test data. The plots present the cumulative distributions for each of the first three markers. Top: before calibration. Bottom: after calibration.

Figure 3 demonstrates the quality of the calibration for individual markers. To analyze whether higher order structures are also preserved during the calibration process we computed the pairwise correlation matrices *C_A_*, *C_B_* for each batch of the test data, before and after calibration. We then computed the difference matrix *C_A_* − *C_B_*. While the Frobenius norm of the difference matrix was 2.82 before calibration, it is 1.11 after calibration, which implies that the calibration process makes the pairwise marker structure between batches significantly more similar. To complete this view, Figure 4 presents the coefficients of the difference matrices before and after calibration, where we can observe that the differences are much smaller after the calibration.

**Figure 4:**
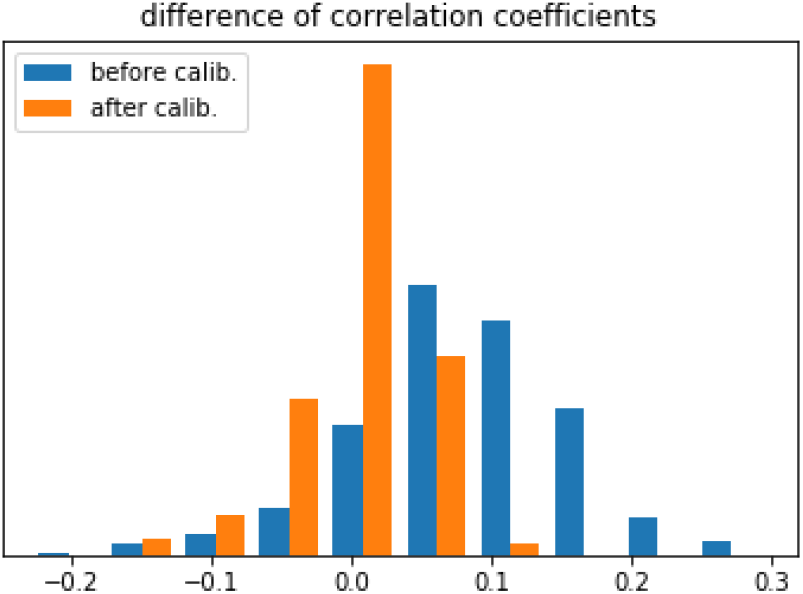
Differences between pairwise correlation coefficients of CyTOF test data before and after calibration. The differences are much smaller after calibration, implying that the correlation structure of the two batches is significantly more similar than before calibration.

Next, we perform quantitative evaluation, by computing the MMD between the source and target batches before and after calibration. The results are shown on Table 1. Each value was computed based on a random subset of size 1000 from each sample; the presented values are mean± std based on three runs. As we can see, calibration decreases the MMD between the distributions of the two batches, on both the train and test data. As one may expect, the MMD on the test data is slightly higher than the train data. In [17] it was shown that MMD-ResNet performs significantly better, in terms of MMD, than popular linear methods for batch effect removal. Although comparison of the current approach to MMD-ResNet is beyond the scope of this manuscript, we remark that it is reported in [17] that the MMD between the source and target batches, *when training a MMD-ResNet on patient 1* was 0.27. Here the corresponding value is 0.26, despite the fact that our system was not trained on this patient, but rather on patient 2, while the data of patient 1 was used merely for testing.

**Table 1:** MMD between the two batches before and after calibration.

| data | before calibration | after calibration |
| --- | --- | --- |
| Train | $.19 \pm .001$ | $.17 \pm .001$ |
| Test | $.26 \pm .01$ | $.19 \pm .01$ |

Finally, to investigate further the preservation of biological structures on the test data, we take a similar approach to [17] and visually inspect the quality of the calibration on the sub-population of Killer T-cells in the 2D subspace of the markers CD28 and GzB, which is shown in Figure 5. As can be seen, the distributions of the Killer T-cells sub-population in the two batches are much closer after calibration.

**Figure 5:**
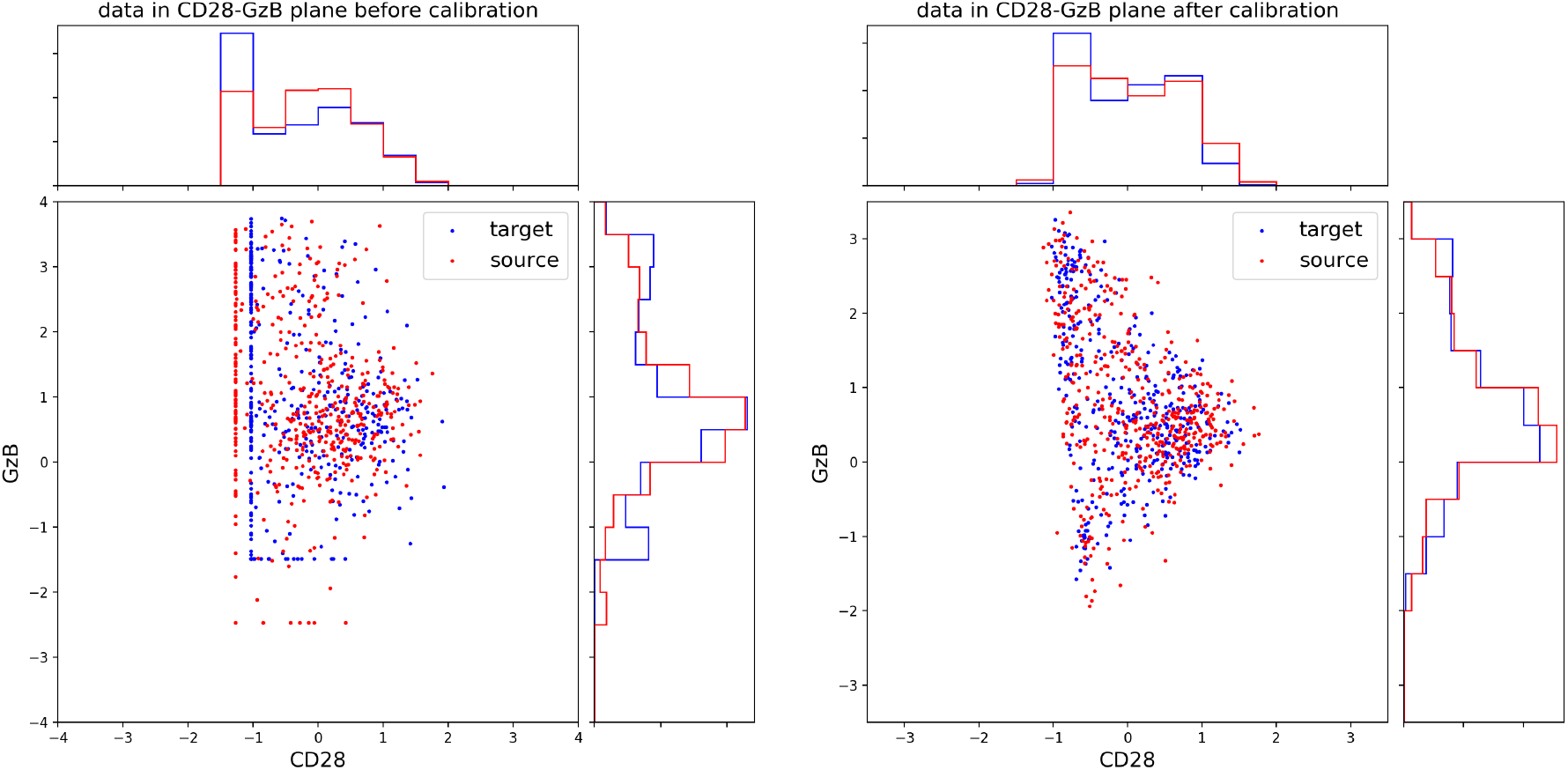
Calibration of CyTOF test data: Killer T-cells in the 2D subspace of the markers CD28 and GzB.

To conclude the CyTOF experiments, we investigated the reconstruction errors and calibration differences through various points of view. In addition to performing high quality calibration on the train data, we also observed similar performance on test data, i.e., on a second patient, having unique biological conditions, whose data was not used during training.

### 4.2 scRNA-seq

Drop-seq [14] is a novel technique for simultaneous measurement of single-cell mRNA expression levels of all genes of numerous individual cells. As a single run of drop-seq typically does not contain enough cells to perform an analysis, multiple runs need to be conducted, a process that might introduce batch effects into the measurements.

In this section we experiment with the publicly available scRNA-seq data described in [17], where a dataset with 13,166 genes is normalized and projected onto the subspace of leading 37 principal components. Altogether, the data contains 27,499 cells in two batches. For a more detailed description of the data we refer the reader to section 4.3 in [17].

Figure 6 shows a T-SNE embedding of data before and after removing the batch effects. As can be seen, our system seems to correctly calibrate the data and align is different sub-populations.

**Figure 6:**
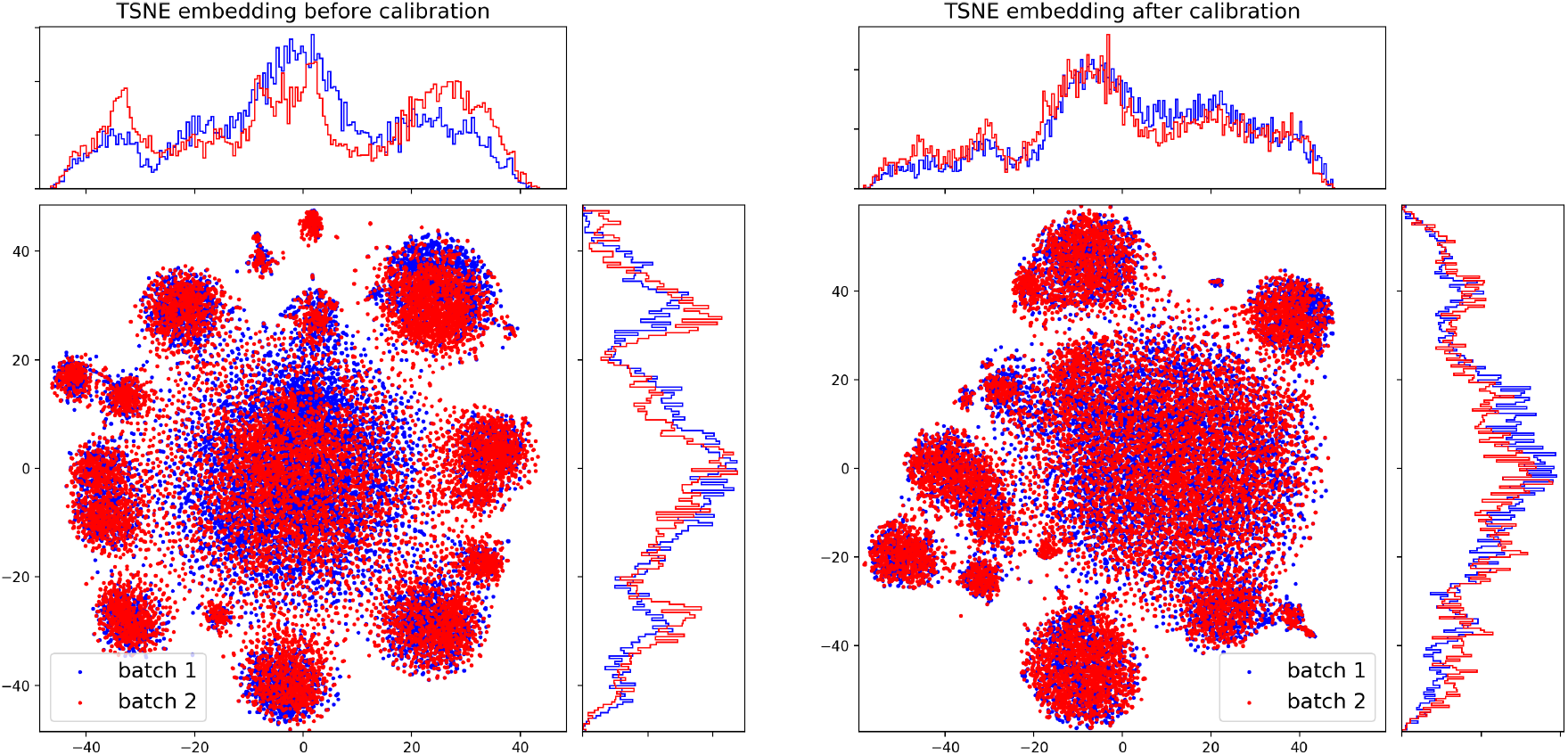
Calibration of scRNA-seq data.

### 4.3 Technical details

All reported results were obtained where *E*, *D_A_*, *D_B_*, Disc all had a basic Multi Layer Perceptron (MLP) architecture, having two hidden layers, each of 20 leaky ReLU units. The code space dimension was 15 in the CyTOF experiments and 20 for the scRNA-seq experiments. We used Adam optimizer with learning rate of 10^−3^ and batch size of 64. Z-transform was applied to all data prior to training. Our publicly available code and data reproduce all reported results.

## 5 Conclusion

We proposed a deep learning-based approach for batch effect removal. Our approach is based on utilizing adversarial loss in order to obtain a encoding of the data which correspond solely to the intrinsic biological state of a subject, along with requiring good reconstruction of the data, which implies that no significant biological information is lost during the calibration process. We demonstrated the performance of our proposed approach on two novel high throughput technologies, CyTOF and scRNA-seq. Moreover, we also demonstrated that our approach can achieve good performance on test data obtained from subjects who are different from the subjects whose data used for training.

## Acknowledgements

The author was partially supported by NIH grant #1R01HG008383-01A1 (PI: Yuval Kluger).

## References

[1] Matthew Amodio and Smita Krishnaswamy. Magan: Aligning biological manifolds. arXiv preprint arXiv:1803.00385, 2018.

[2] Matthew Amodio, Ruth Montgomery, Jenna Pappalardo, David Hafler, and Smita Krishnaswamy. Neuron interference: Evidence-based batch effect removal. arXiv preprint arXiv:1805.12198, 2018.

[3] Matthew Amodio, Krishnan Srinivasan, David van Dijk, Hussein Mohsen, Kristina Yim, Rebecca Muhle, Kevin R Moon, Susan Kaech, Ryan Sowell, Ruth Montgomery, et al. Exploring single-cell data with multitasking deep neural networks. bioRxiv, page 237065, 2017.

[4] Martin Arjovsky, Soumith Chintala, and Léon Bottou. Wasserstein gan. arXiv preprint arXiv:1701.07875, 2017.

[5] Ian Goodfellow, Jean Pouget-Abadie, Mehdi Mirza, Bing Xu, David Warde-Farley, Sherjil Ozair, Aaron Courville, and Yoshua Bengio. Generative adversarial nets. In Advances in neural information processing systems, pages 2672–2680, 2014.

[6] Arthur Gretton, Karsten M Borgwardt, Malte J Rasch, Bernhard Schölkopf, and Alexander Smola. A kernel two-sample test. Journal of Machine Learning Research, 13(Mar):723–773, 2012.

[7] Ishaan Gulrajani, Faruk Ahmed, Martin Arjovsky, Vincent Dumoulin, and Aaron C Courville. Improved training of wasserstein gans. In Advances in Neural Information Processing Systems, pages 5767–5777, 2017.

[8] Kaiming He, Xiangyu Zhang, Shaoqing Ren, and Jian Sun. Deep residual learning for image recognition. In Proceedings of the IEEE conference on computer vision and pattern recognition, pages 770–778, 2016.

[9] Irina Higgins, Loic Matthey, Arka Pal, Christopher Burgess, Xavier Glorot, Matthew Botvinick, Shakir Mohamed, and Alexander Lerchner. beta-vae: Learning basic visual concepts with a constrained variational framework. 2016.

[10] Diederik P Kingma and Max Welling. Auto-encoding variational bayes. arXiv preprint arXiv:1312.6114, 2013.

[11] Jeffrey T Leek, Robert B Scharpf, Héctor Corrada Bravo, David Simcha, Benjamin Langmead, W Evan Johnson, Donald Geman, Keith Baggerly, and Rafael A Irizarry. Tackling the widespread and critical impact of batch effects in high-throughput data. Nature Reviews Genetics, 11(10):733, 2010.

[12] Huamin Li, Uri Shaham, Kelly P Stanton, Yi Yao, Ruth R Montgomery, and Yuval Kluger. Gating mass cytometry data by deep learning. Bioinformatics, 33(21):3423–3430, 2017.

[13] Mingsheng Long, Yue Cao, Jianmin Wang, and Michael I Jordan. Learning transferable features with deep adaptation networks. arXiv preprint arXiv:1502.02791, 2015.

[14] Evan Z Macosko, Anindita Basu, Rahul Satija, James Nemesh, Karthik Shekhar, Melissa Goldman, Itay Tirosh, Allison R Bialas, Nolan Kamitaki, Emily M Martersteck, et al. Highly parallel genome-wide expression profiling of individual cells using nanoliter droplets. Cell, 161(5):1202–1214, 2015.

[15] Noam Mor, Lior Wolf, Adam Polyak, and Yaniv Taigman. A universal music translation network. arXiv preprint arXiv:1805.07848, 2018.

[16] Anthony Rios, Ramakanth Kavuluru, and Zhiyong Lu. Generalizing biomedical relation classification with neural adversarial domain adaptation. Bioinformatics, 1:9, 2018.

[17] Uri Shaham, Kelly P Stanton, Jun Zhao, Huamin Li, Khadir Raddassi, Ruth Montgomery, and Yuval Kluger. Removal of batch effects using distribution-matching residual networks. Bioinformatics, 33(16):2539–2546, 2017.

[18] Matthew H Spitzer and Garry P Nolan. Mass cytometry: single cells, many features. Cell, 165(4):780–791, 2016.

[19] Gil Tabak, Minjie Fan, Samuel J Yang, Stephan Hoyer, and Geoff Davis. Correcting nuisance variation using wasserstein distance. arXiv preprint arXiv:1711.00882, 2017.

